# Modeling Particle Transport In Biomedical Flows Using Implicit Geometry Representations

**DOI:** 10.64898/2026.06.07.730719

**Authors:** J. Scott Malloy, Sreeparna Majee, Akshita Sahni, Ricardo Roopnarinesingh, Aditya Balu, Adarsh Krishnamurthy, Debanjan Mukherjee

## Abstract

Computational analysis of physiological and biomedical systems necessitate efficient geometry representations for high fidelity model predictions, including patient or device specificity. Particle-based Lagrangian computational approaches comprise a valuable approach to gain insights from quantitative velocity and pressure data from computational models. Examples include particle dynamics and transport in human vasculature for diseases such as stroke, thrombosis, and embolisms; and modern targeted drug delivery systems in the vascular network and respiratory airways. However, current particle simulation approaches can bear significant computational expense that scales with both number of particles and background fluid mesh resolution. A significant determinant of this computational expense is the contact resolution between particles and anatomically realistic vessel wall. Here, we develop an efficient particle dynamics model that leverages an implicit representation of real anatomical features using a signed distance field to efficiently resolve particle-wall contact. We outline the underlying algorithmic details, followed by a systematic illustration of performance and accuracy using simplified and analytically defined geometries and flow fields. Subsequently, we present a representative simulation of embolic particles along a human vascular segment where we compare our distance field-based approach against classical wall-contact checks based on assessing particle boundary intersection with triangulated surface mesh. Our approach transforms the underlying Lagrangian contact detection operation into an equivalent Eulerian operation, significantly speeding up bulk particle dynamics computations without significantly impacting accuracy or geometric fidelity.

## 1 Introduction

Geometric complexities are a central feature in physiological and biomedical systems. Examples include complex shapes and morphologies of structures for biomedical devices and implants, and anatomical complexities for various regions of interest in the human body. Analysis of fluid flow, mass transport, and fluid structure interactions, remain central to a wide range of applications involving physiological and biomedical systems. For appropriate fidelity, such computational analyses must be geometry aware or anatomy aware [1–4]. This motivates the need for innovative avenues to incorporate flexible high-fidelity geometric representations to high-resolution computational analysis workflows. Here, we focus on one such key modality: *robust Lagrangian analysis workflows for biomedical and physiological flow and transport phenomena assessment*. Lagrangian computations broadly refer to mesh-free particle-based numerical approaches, where the trajectories of either massless idealized tracer particles or finite-size finite-inertia physical particles are computed numerically within a given fluid flow. While standard computational fluid dynamics analysis provides detailed information about velocity and pressure fields, often discerning physiologically relevant insights on flow and transport requires quantitative analysis that cannot be fully illustrated using only the velocity and pressure data. Lagrangian analysis provides a robust alternative [5, 6], enabling information on critical quantities like residence time for quantifying flow stasis [7], hidden coherent structures in nonlinear biomedical flows that organize mass transport [8–11], background stress and strain topology in the fluid flow [12], quantification of stress or strain exposure for platelets and erythrocytes for estimation of platelet activation and blood damage [13, 14], and also transport of actual particulate species for particle-based drug delivery [15–17] or fragmented clot movement in the vascular network [18, 19]. More broadly particle-based methods can also include mesh-free approaches where discrete particles comprise computational units to compose discrete mathematics needed for simulating flow or mechanics. Examples of such techniques include, but are not limited to, smoothed particle hydrodynamics [20], discrete element methods [21], dissipative particle dynamics methods [22], and particle finite element method [23].

For such particle-based methods, accuracy relies heavily on robust resolution of particle interaction with the boundary walls of the domain. For vascular and respiratory applications, this will comprise particle interactions with vessel (or airway) walls. This component of the computation can be resource intensive, and in many instances, a significant fraction of computational cost is attributed to particle-wall contact resolution. The cost is influenced by the aspect of geometric complexity stated above, with common Lagrangian computations requiring checks of a large number of particles with anatomically realistic vessel walls that are arbitrarily curved and tortuous, and can vary substantially in cross-sectional area. Inaccuracies in particle-wall contact and subsequent particle motion updates can lead to inaccuracies in near-wall particle distribution, particle trajectories, and associated quantitative metrics. Hence, there are several existing approaches that address this computational need, with algorithms that range from naïve particle contact checks with each triangle element of a surface mesh [24] using geometric or analytical relations to efficient spatial partitioning of the wall boundary mesh and tree-based algorithms [25] to reduce computational complexity of the candidate particle-wall contact pair checking. In some implementations, the walls themselves are often reconstructed as particles for simplified contact checking. Space partitioning approaches are commonly implemented in existing state-of-art tools (such as, for instance, the VTK library, which is employed for some of the analyses in this study). These existing approaches face challenges when resolving the specific geometric complexities of anatomical structures such as a large heart to brain arterial network, or the interstices of an anatomically realistic bone scaffold specimen. Despite the availability of existing tools, therefore, there remains a need for establishing alternative methodologies that can enable tackling this underlying geometric complexity for robust particle-wall contact resolution in Lagrangian computations (and other relevant computational analysis).

Motivated by this research question, here we outline a particle-wall interaction modeling approach using a geometric entity named the Signed Distance Field (SDF) for implicit representation of anatomical features. SDF have been used extensively for a wide range of applications, such as shape manipulation [26], shape and volume representation [27], constructive solid geometry (CSG) Boolean operations [28], and interaction between geometric entities [29–31]. For any given geometry, SDF represents a spatial function where the function value at any point is its distance to the closest boundary point. Our approach specifically leverages this fact, and known mathematical properties of the SDF, to develop a novel transformation of the Lagrangian contact check with the wall into a fast Eulerian interpolation of the distance field that dictates particle proximity to the wall. We leverage a GPU-accelerated algorithm for generating fast-SDF estimates, demonstrated in prior works, which can represent the anatomical structures such as vascular trees with high resolution. As an implicit field-based representation of the geometry, this approach enables flexible incorporation of complex geometries; while providing gains in computational efficiency and accuracy of wall-interaction predictions. In this manuscript, we outline the algorithm and focus on establishing three key aspects: (1) the accuracy of our approach using simplified models based on analytical flows; (2) the computational speedup using our approach based on an idealized model problem; and (3) demonstration of our methodology in a realistic biomedical application which involves the transport of thrombo-embolic particles within patient-specific models of vascular segments in the brain.

## 2 Particle Transport Framework

### 2.1 Generation of Implicit Geometry Representation

We represent our computational domain for the physiological or biomedical system of interest (*for instance, a vascular segment*) as Ω, with a clearly delineated boundary *∂*Ω without any holes. The Signed Distance Field is then represented mathematically as:

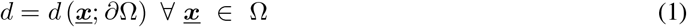

The zero level-set of this field is the wall or boundary of the domain. The distance *d* is therefore maximum at any cross-section at the geometric midpoint or center of that cross-section. For internal flow domains such as a blood vessel, the maximum values of *d* will therefore follow the centerline of the structure (*information that can be used to directly extract the centerline as a geometric object, or to further use it for skeletonization of the vascular structure for reduced-order network representations*). The gradient of the SDF evaluated at the wall points along the steepest descent of distance, and at the boundary surface *∂*Ω it is equivalent to the inward surface normal:

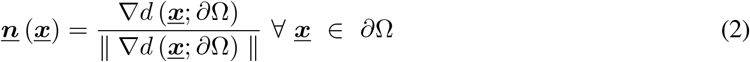

Furthermore, the Hessian of the distance field *H*(***<u>x</u>***) = ∇∇*d* denotes the curvature of the geometry domain. We compute the distance transform on a regular voxel grid using the GPU-accelerated hybrid marching wavefront scheme of Balu *et al*. [32]. Distance transforms are discrete representations of distance fields on a regular voxel grid; two principal limitations of such representations are their computational expense and the discretization errors introduced by the object representation. The algorithm proceeds in two steps: (i) computing the distance values and their theoretical bounds for boundary voxels, and then (ii) propagating those values to the remaining voxels. Boundary voxels are defined as those voxels in the grid that have at least one neighbor with a differing occupancy indicator value. In 3D, the neighborhood is taken as all 26 voxels sharing a face, edge, or vertex with the current voxel. The minimum distance from the object boundary to the voxel center for each boundary voxel is computed from the boundary representation; for a voxel-based representation, the theoretical bounds on this distance are 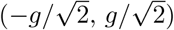, where *g* is the voxel grid spacing. Boundary voxels are assigned a nominal distance value of zero. Once boundary voxel distances are established, distances are propagated outward by one neighborhood ring of voxels per pass. In 2D, this marching can proceed in orthogonal and diagonal directions; the same principle extends to 3D with additional tridiagonal contributions [32].

A fundamental disadvantage of standard marching approaches is that directly propagating distance values accumulates error across successive passes: diagonal marches of length 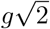 composed from orthogonal steps introduce a systematic overestimate, reaching up to approximately 7% relative error in 2D and 12.8% in 3D. The hybrid approach of Balu *et al*. [32] eliminates this accumulation by storing and propagating the index of the closest boundary voxel at each grid point rather than the distance value itself. The true Euclidean distance to that boundary voxel is then recomputed exactly at each update, such that the only remaining error in the final distance transform arises from the tessellation of the surface geometry rather than from the marching procedure. To handle the race conditions that arise from parallelizing this update across the GPU, a ping-pong scheme is employed: two copies of the distance field matrix are maintained, with the previous pass (DTM-A) used as the reference for all comparisons and the updated values written to the current pass (DTM-B), after which the two are swapped. The algorithm terminates when the total sum of distance field values does not change between two successive iterations, or when a maximum of 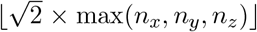 marching passes have been completed, a bound that follows from the maximum Euclidean distance in the grid being traversable in that many steps of minimum size *g*. The resulting SDF is then interpolated onto the nodes of the computational domain mesh for subsequent particle proximity and contact detection, as described in Section 2.2.

### 2.2 Particle Collision Algorithm

The proposed wall contact detection methodology is conceptually illustrated in Figure 1. Specifically, once the SDF is generated as an implicit representation of the closest point projection on a voxel-form of the computational domain of interest, the computed SDF field *d* (<u>***x***</u>; *∂*Ω) is re-interpolated onto the nodes of the mesh of the computational domain; which can be an unstructured grid with a specified mesh resolution. As an individual particle trajectory is integrated from one time-step to the next, the SDF *d* (<u>***x***</u>; *∂*Ω) is interpolated onto the particle centroid location along with other flow variables (e.g. velocity, pressure). This calculation relies on identifying the cell/element that contains the particle centroid. For a structured grid, this can be identified using simple binning and indexing algebra. For an unstructured grid on the other hand, the element can be identified using tree-based search algorithms (*such as the vtkCellTreeLocator implementation in the VTK library*), or a geometric cell-crawling algorithm based on barycentric coordinates for tetrahedral elements as outlined in [33]. While the interpolated velocity fields, for instance, are used to subsequently update particle positions for the next time-step, the interpolated SDF field is used for resolving particle-wall interactions. Additionally, the SDF gradient is used to identify the inward facing wall normals; and this leads to definitions of the wall normal and wall tangential components of the particle velocities as stated below:

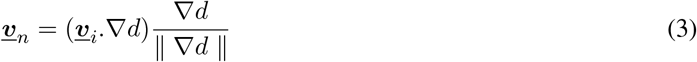

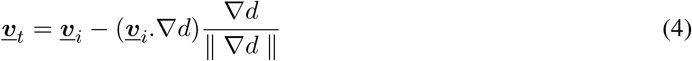

Based on these details, for a finite-size particle approaching a surface *∂*(Ω), contact is identified to have occurred, if the interpolated SDF at the particle centroid location is less than or equal to the particle radius (*assuming spherical particles for the purpose of this description*).

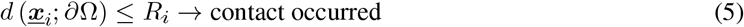

where, ***<u>x</u>***_*i*_ represents the current position, and *R*_*i*_ represents the particle radius for the *i*’th particle respectively. For massless point particles or Lagrangian tracers, the above Eq. 5 is modified by replacing *R*_*i*_ using a numerically small penalty *ϵ* ≈ 1*e* − 10 − 1*e* − v14 for resolving wall interactions. Furthermore, in the event a contact or collision is detected, the particle velocity and/or position is subsequently updated using SDF field information. Specifically, velocity is updated along the normal and tangential direction as estimated based on interpolated SDF fields, using effective restitution coefficients. The normal velocity update then can be computed as follows:

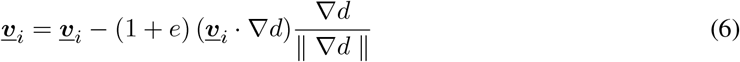

**Figure 1:**
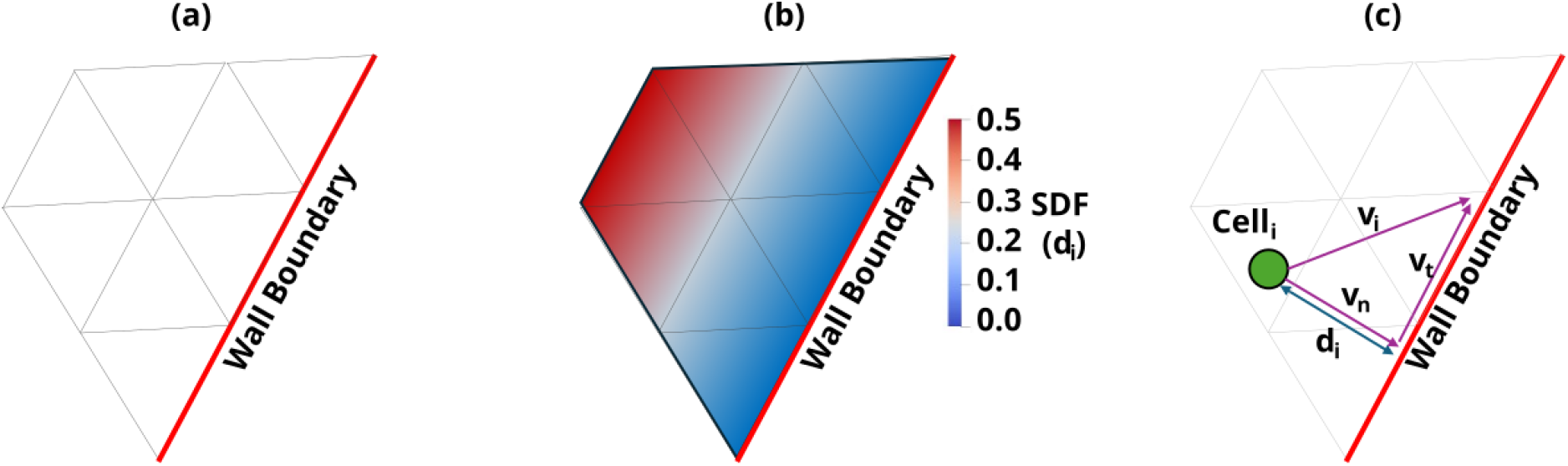
Proposed Eulerian contact detection algorithm, starting from a representative STL model (a) with the computed SDF field showing the distance to the nearest wall (b), and a particle coming in proximity with the wall with velocity tangential and normal to the wall (c)

We note that the above formulation uses a constant valued restitution. When an immersed particle happens to collide with the wall, a viscous lubrication fluid layer is formed, increasing pressure loading on the particle and thus deforms it [34, 35]. This mechanism adds more dissipation on the contact dynamics, causing effective restitution to change, which can be modeled as a velocity dependent restitution coefficient, devised for biomedical fluid flows in prior works [18]. This can be directly incorporated in Eq. 6. In addition, the tangent and bi-tangent directions can be further resolved by noting that the local maxima at any point along a vessel segment will align with the centerline of the vessel, and derivative of the centerline yields the tangent along the axial direction. The cross-product of the normal and tangent yields the bi-tangent direction. This further enables writing velocity update equations equivalent to Eq. 6 for the tangential components of the velocity. The complete algorithmic implementation is outlined in Algorithm 1.

## 3 Framework Evaluation in a Model Tortuous Vessel

One key geometric complexity of human vascular anatomy lies in variable curvature and tortuous vessel segments, which can lead to complex wall-interaction scenarios for particle and Lagrangian computations. Here, we demonstrate the application of our proposed SDF-based collision algorithm in a 3D meshed domain comprising a mock curved or serpentine vessel, illustrated in Figure 2, panel a; an idealization of geometries common in anatomical and microfluidic systems. The experiments with this setup were intended to assess the algorithm’s scaling costs in a collision rich environment within a representative internal flow domain with varying curvature, analogous to vascular segments. The geometry was created using Gmsh. Once created, the SDF was computed and mapped onto the vessel domain mesh comprising linear tetrahedral elements; but without any fluid flow velocity data computed on the mesh (*making this a purely geometric contact evaluation experiment*). The computed SDF is shown in Figure 2, panel b., where the SDF value is maximum along the centerlines at 0.1 m (*which is the radius of the tube geometry*). The domain was meshed at 8 varying resolutions, for comparison of computational cost for SDF-based collision checks. The mesh sizes and the number of associated wall surface mesh elements have been listed in Table 1.

**Table 1:** Mesh refinement levels showing the number of volume and wall surface elements for numerical experiments with the sample curved vessel geometry discussed in Section 3.

| Mesh Number | Volume Elements | Wall Surface Elements |
| --- | --- | --- |
| 1 | 7,407 | 2,124 |
| 2 | 18,610 | 4,286 |
| 3 | 33,544 | 6,778 |
| 4 | 80,300 | 12,710 |
| 5 | 178,526 | 22,502 |
| 6 | 560,688 | 50,234 |
| 7 | 1,304,434 | 89,230 |
| 8 | 4,194,316 | 199,669 |

**Figure 2:**
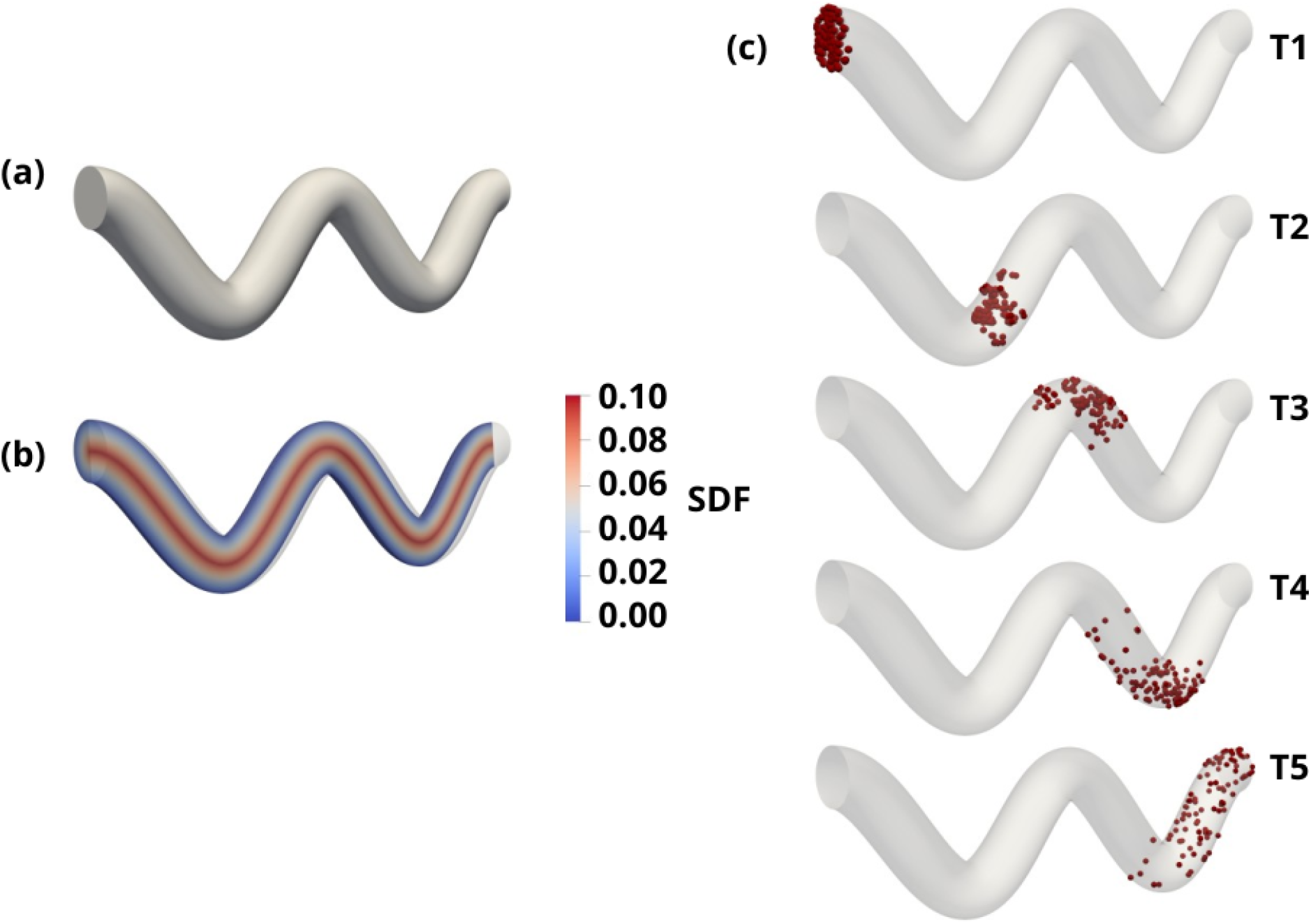
Configuration of mock tortuous vessel including (a) raw geometry, (b) generated SDF field with maximum value in centerline equal to radius of cross-section and (c) sample of particle positions for SDF-algorithm tracked particles at 5 time steps

An ensemble of particles were randomly seeded at the inlet of this geometry, with initial velocities of 1.0m/s pointed normal to the inlet cap and 0.5m/s tangential to the cap, to align their initial trajectories with the shape of the vessel. Each particle was subsequently tracked for 4.0 seconds at a time step of 0.001 second for a total of 4000 time steps. Particle ensemble sizes ranged from 50 to 10000 across the varying mesh resolutions outlined above. Given that the background mesh has no fluid flow data, the trajectory of each particle will be solely due to their initial velocity and any changes resulting from wall collisions, enabling a controlled analysis of SDF-based collision resolution efficiency. Figure 2, panel c. illustrates the particles moving through this domain, due to wall collisions, for 5 snapshots at successive instants in time.

The overall computational time to resolve 4.0 seconds of the ensemble dynamics was estimated across all the particle ensemble sizes and all the mesh resolutions; spanning multiple collisions for each simulation that was resolved using the SDF-based approach. The results are illustrated in Figure 3, where panel a. depicts the compute time across varying mesh resolution, for each ensemble; and panel b. depicts the compute time across varying ensemble sizes, for each mesh resolution. The data clearly illustrates that while compute times scale directly linearly with the number of particles, they remain essentially flat with mesh resolution showing significantly less variation over the large mesh resolution range considered here. We note that, considering the number of particles to be *N*_*p*_, and the mesh resolution represented in terms of number of simplices in the wall surface mesh *N*_*s*_, the naive complexity estimate will be 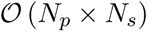. Advanced mesh-partitioning or tree-based search algorithms can further reduce the dependency on mesh resolution here to 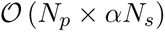 with *α* being a scaling factor less than unity. Our analysis here depicts that with the SDF based approach we can potentially reach a complexity of 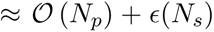; meaning that leading order complexity is independent of mesh resolution (*even if total cost shows weak dependence on mesh resolution*). This demonstrates the benefit of reducing the wall collision check to a purely Eulerian operation, in a setting with more complex geometry that yields more collisions in comparison to the cylinder model.

**Figure 3:**
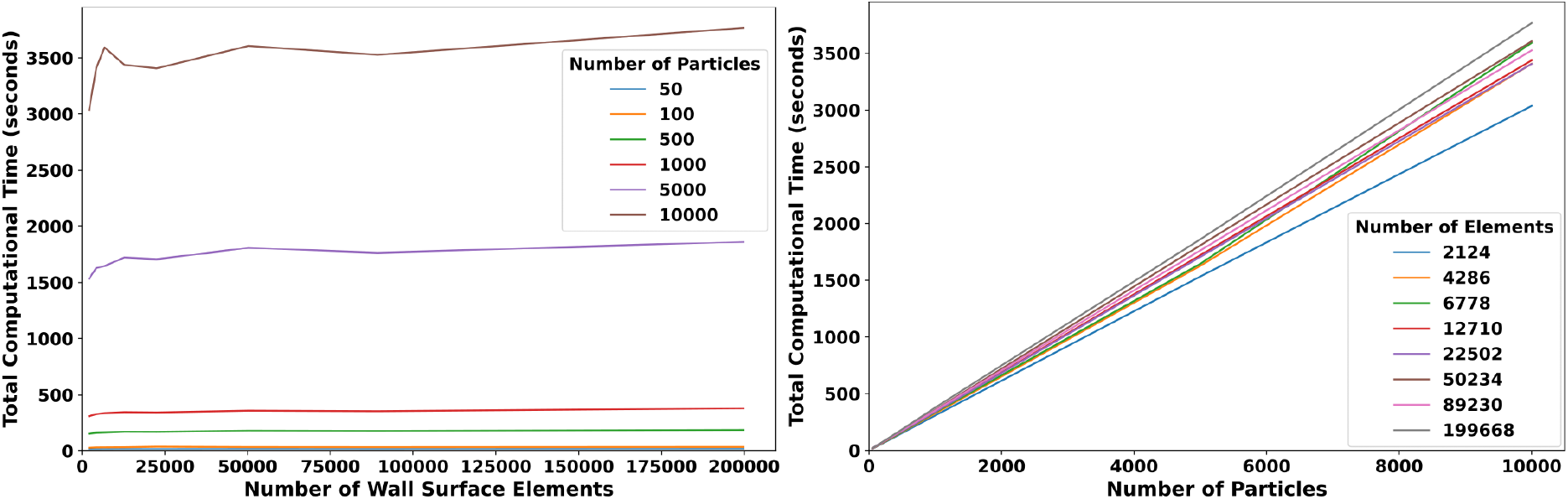
Total computational time scaling for SDF evaluation: (a) increasing mesh resolution and (b) increasing number of particles tracked.

## 4 Framework Validation Using Analytical Solutions

Here, we outline a series of numerical experiments that illustrate the numerical accuracy of the SDF-based approach in conjunction with computational cost for particle movement within a cylindrical tube with a circular cross-section, with an imposed parabolic fluid flow velocity profile along the axial direction. We used a cylinder of length *l* = 2.0*m* axially aligned along *x*-direction, and radius *r*_*cyl*_ = 0.5*m* (*crosssection along y* −*z plane*), shown in Figure 4, panel a. Given the cylinder shape, proximity of a particle to the wall was computed analytically for controlled comparison against SDF-based particle checks. The geometry was meshed at varying resolutions (similar to Section 3), and computed SDF was mapped onto the mesh. The mesh sizes, and number of surface elements are shown in Table 2. A parabolic background axial velocity was prescribed within the cylinder with a maximum velocity of 0.2 m/s. Within this flow, trajectories of particle ensembles of varying sample sizes from 50 to 5000, were numerically integrated. For each ensemble, particle radius was set at *R*_*p*_ = 0.25 mm, and the particles were initially seeded randomly at the cylinder inlet, and these initial positions were held fixed between two simulation cases: one with an SDF-based collision check simulation, and the other with an analytically determined collision check. The particles were assigned an initial velocity of 0.2 m/s along the axial direction, and 2.0 m/s along the tangent to the inlet cap; ensuring that each particle would collide with the wall at least twice. Each particle position and velocity was computed over time based on Stokes drag due to the background flow, and velocity updates from wall contacts. These particles were tracked for 4.0s to allow particles to reach the outlet at a time step of 0.001s for a total of 4000 time steps. The underlying differential equation describing the particle dynamics is as follows:

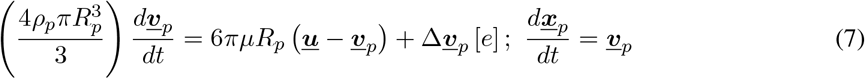

with fluid dynamic viscosity *µ* = 0.0004 kg/mm.s, particle density *ρ*_*p*_ = 0.11 kg/mm^3^, which together yield a momentum response time ≈ 4.0 s; enabling an appropriately intermediate regime where the particle velocity does not fully align with fluid flow within the computation horizon. The term Δ***<u>v</u>***_*p*_[*e*] indicates the updated velocity due to wall contact, as a function of the restitution coefficient *e*. The simulations with the SDF-based algorithm involved a computed update for the particle positions, followed by an SDF-based check for collisions, and computing the restitution-based velocity update when collisions occurred. The simulations with analytical check simply replaced the wall-contact resolution with an analytical distance and normal calculation. The distance to the wall was computed by 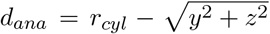 with (*y, z*) being the radial position of the particle. Contact was resolved for a velocity update when *d*_*ana*_ ≤ *R*_*p*_ where *R*_*p*_ denotes the particle radius. The normal vector was computed from the location of the particle relative to the center of the cylinder: 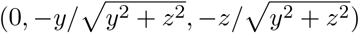.

**Table 2:** Mesh refinement levels showing the number of volume and wall surface elements.

| Mesh Number | Volume Elements | Wall Surface Elements |
| --- | --- | --- |
| 1 | 42,467 | 4,376 |
| 2 | 70,453 | 6,240 |
| 3 | 134,310 | 9,782 |
| 4 | 303,829 | 17,040 |
| 5 | 958,628 | 38,462 |
| 6 | 2,289,889 | 67,828 |
| 7 | 7,368,872 | 151,958 |

**Figure 4:**
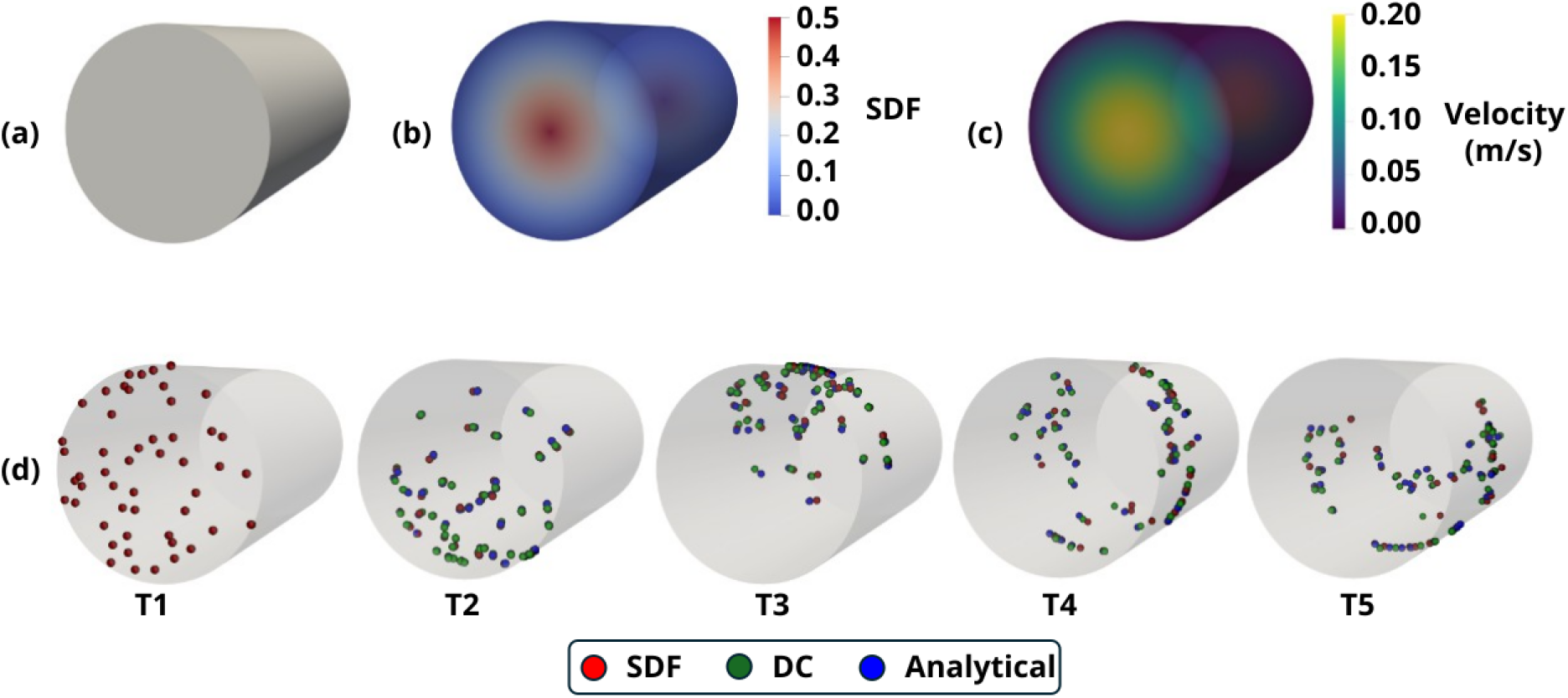
Configuration of model used for validation including raw cylinder geometry (a), generated SDF field with maximum value in centerline equal to radius of cylinder (b) and sample of particle positions for SDF-algorithm (red) Direct Check algorithm (green), and analytical (blue) tracked particles at 5 time steps

A non-SDF grid-based particle-wall contact resolution was also implemented for comparison, referred to here as *Direct Check* (DC). This leveraged a conventional cell-partitioning based algorithm as packaged in the vtkStaticCellLocator method implementation in VTK library. The locator identified the wall element nearest to the particle. The closest point on that element was determined, and the vector from the particle to this point was computed and normalized to obtain the surface normal for collision handling. This DC computation was implemented to quantitatively illustrate the time scaling advantages that the SDF algorithm has in comparison to conventional checks without increasing errors in particle trajectory estimates. For both the computations using SDF-based approach and DC-approach, as well as for the analytical computation, the particle positions were computed for each mesh resolution: referred to as ***<u>x</u>***_*SDF,i*_, ***<u>x</u>***_*DC,i*_, and ***<u>x</u>***_*ANA,i*_ respectively for the *i*’th particle in the ensemble. Snapshots of these particle positions at varying instants in time have been illustrated in Figure 4, panel c; showing ***<u>x</u>***_*SDF,i*_, ***<u>x</u>***_*DC,i*_, and ***<u>x</u>***_*ANA,i*_ in red, green, and blue respectively. At each time instance, and for each ensemble, an error metric was computed by comparing the particle positions as follows:

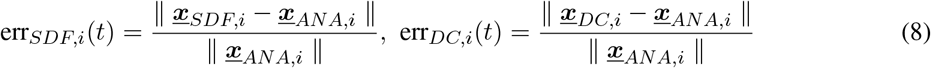

Additionally, the computational time for the trajectories using the SDF-based and the DC algorithm were also recorded for each ensemble size and mesh size. Figure 5 presents the ensemble averaged and time averaged errors stated in Eq. 8, evaluated for the SDF-based calculations and DC calculations, quantified with respect to mesh resolution. Both error metrics decrease with increasing mesh resolution, with an initial sharp decay in error towards a plateau (*at around 1%*) at higher resolutions. Our SDF-based algorithm leverages multiple underlying approximations such as discretization of geometry into a triangular mesh, mapping computed continuous SDF function onto the discretized volume mesh, and locating particles in individual cells. Despite potential errors originating from all these approximations, we note that the overall averaged error behavior remains low and the computations remain stable across mesh resolutions. Figure 5 also shows the corresponding error behavior for the DC-version of the calculations, demonstrating same trend of error reduction with mesh resolution. However, the DC-algorithm based errors remain consistently higher than our proposed SDF-based approach.

**Figure 5:**
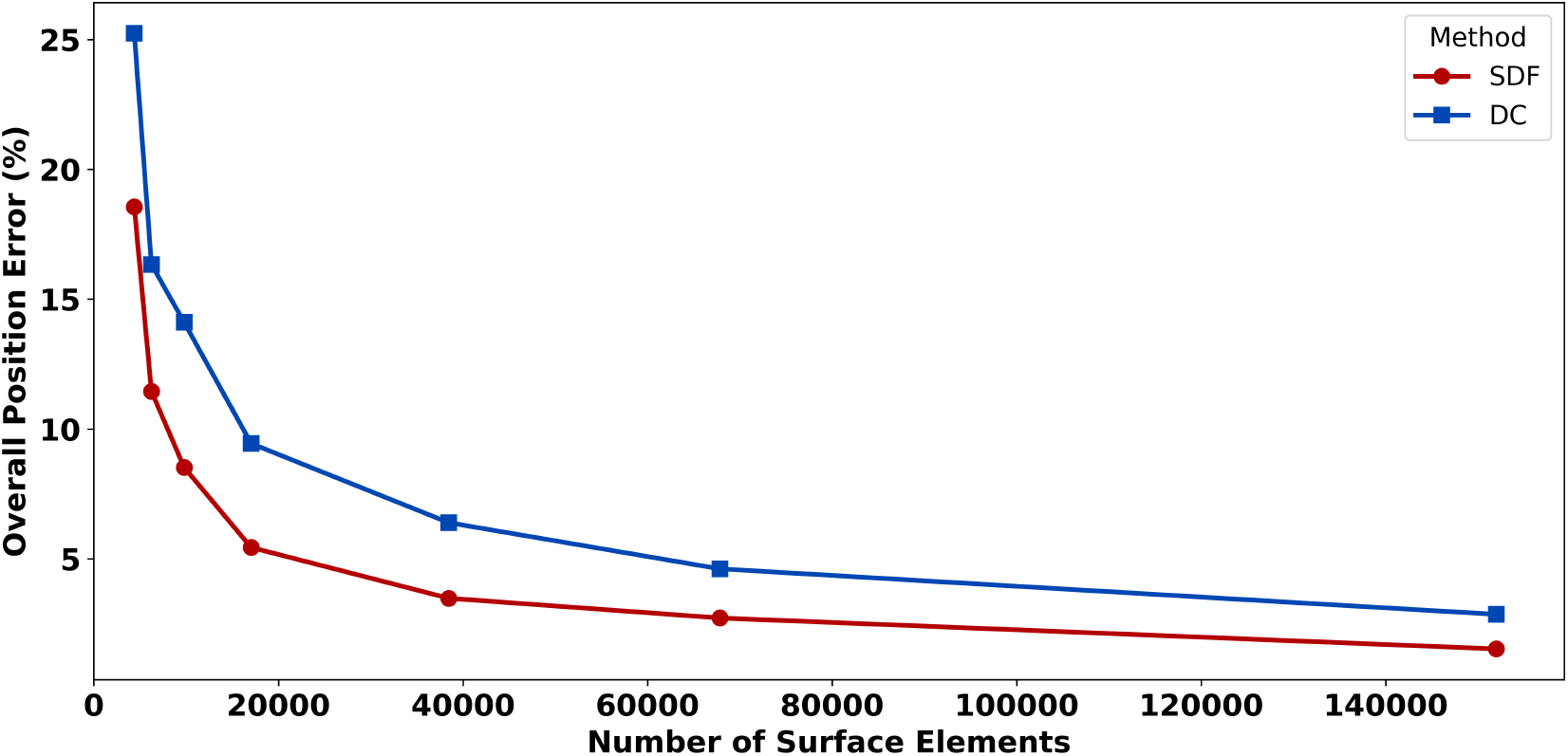
Position error of analytical vs Signed-Distance Field (SDF) and analytical vs Direct Check (DC) computed particle trajectories, averaged over all 6650 particles and time steps.

The results above indicate clearly that at higher mesh resolutions, the SDF field can capture finer geometric details, allowing higher fidelity estimates of the calculated distances between the particle and the wall (*in this case, in reference to the analytical result*). Additionally, for both the SDF-based and DC-based computations, Figure 6 depicts the ensemble averages of the position errors defined in Eq. 8, for each ensemble tracked over time. The errors start from zero, and with time, as collisions occur, the error increases, demonstrating temporal variations that align with timing of the collisions. Specifically, each collision is associated with a temporary dip in the error, as the wall keeps the particles close together. Following a collision, the error is observed to rise (*until the next collision*) as the trajectories keep deviating from the analytical reference post collision. Over time, as the number of collisions become more defuse, the significant bump in the error behavior diminishes, and the position error plateaus. Once again, we observe that the DC-based errors are consistently higher than the SDF-based case.

**Figure 6:**
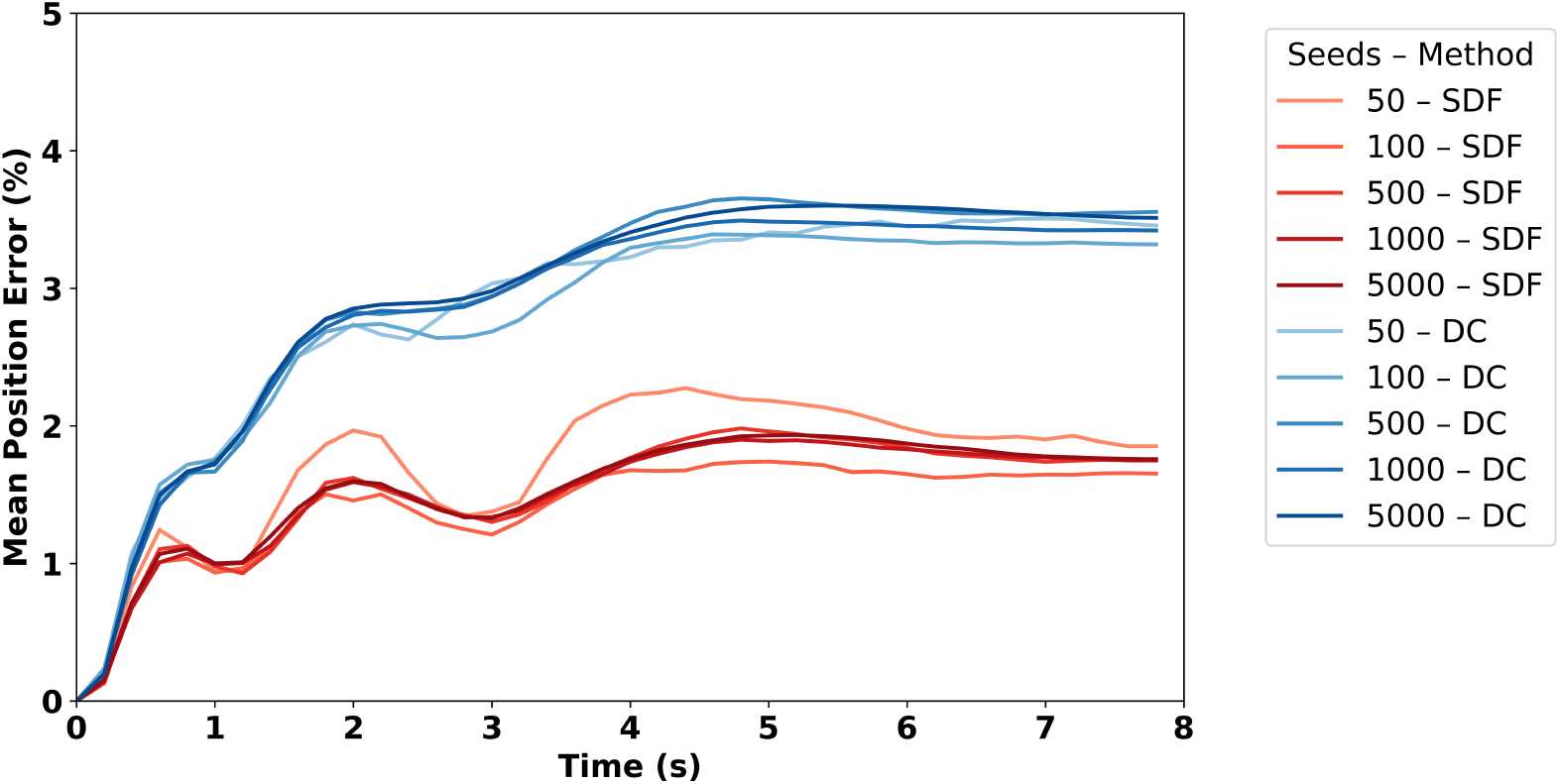
Position error of analytical vs Signed-Distance Field (SDF) and analytical vs Direct Check (DC) computed particle trajectories, for highest mesh resolution (151,958 surface elements), plotted over time

Finally, the total computational cost for the SDF-based and DC-based implementations as function of particle ensemble size and mesh resolution is illustrated in Figure 7a. The observed computational costs for the SDF-based approach demonstrates again a significantly weaker dependency on mesh resolution compared to DC-based approach, and is primarily determined by particle ensemble size, with the computational complexity described as in Section 3. Additionally, for each ensemble, the computational cost for the SDF-based approach is lower than the DC-based approach. Together, this complete analysis reveals that the SDF-based approach is more accurate than equivalent mesh-partitioning type approaches for wall contact (*the common approach in a wide range of particle simulation tools*); while staying computationally cheaper, with computational cost scaling mainly with particle count but weakly with mesh resolution. We note that these trends are for a simplified control case of axial flow within a cylinder, and the trends for accuracy and computational cost as illustrated here are anticipated to also hold as problem complexity increases. This is because, with the SDF check we have converted an essentially Lagrangian operation into an essentially Eulerian grid-based computation. In many applications, this Eulerian operation already occurs to determine other metrics of the cell that the particle is in, such as the velocity within that region, removing additional overhead for the particle calculations. This renders a marked increase in computational efficiency of the SDF implementation in comparison to the conventional wall check algorithms.

**Figure 7:**
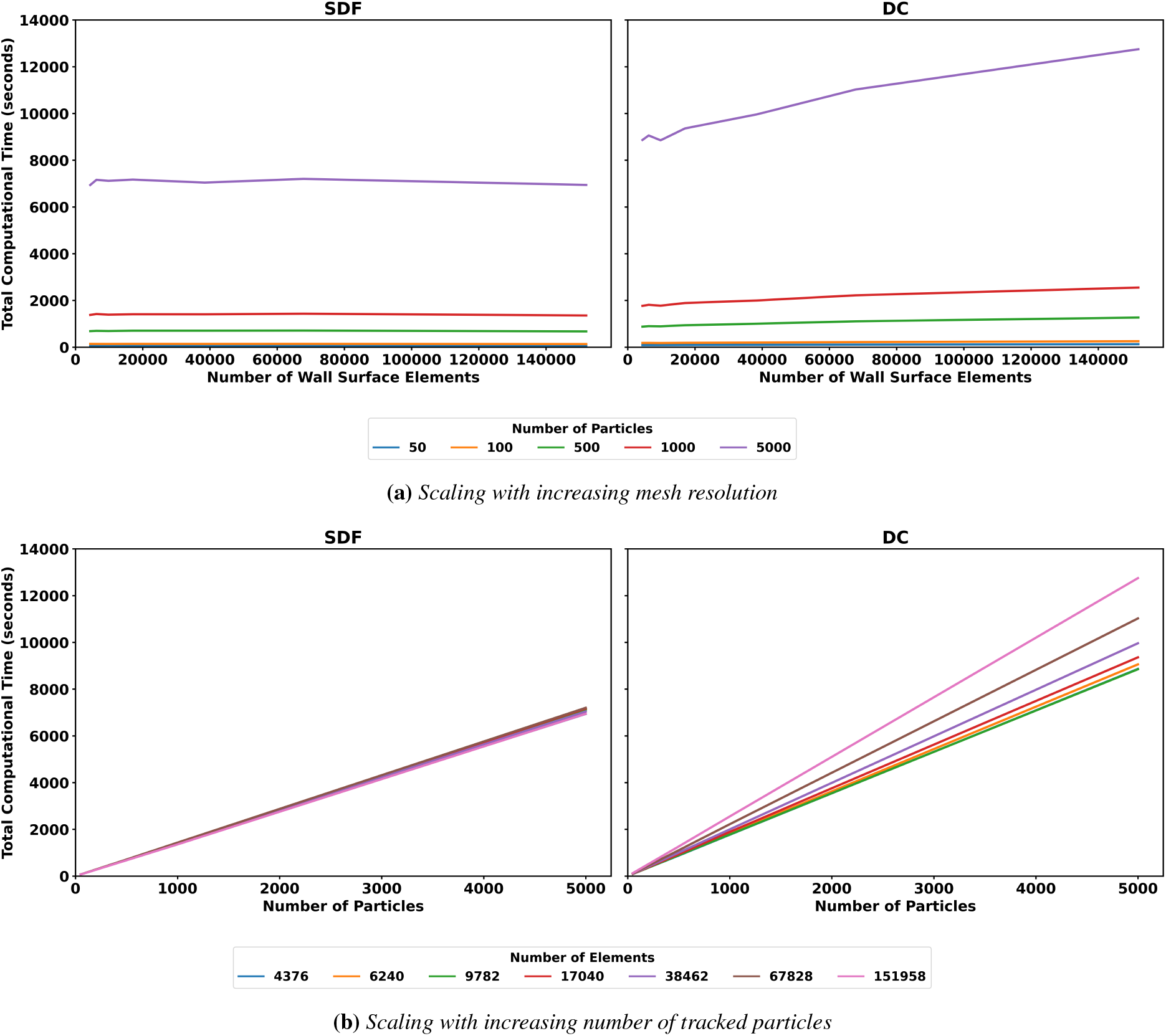
Total computational time scaling for Signed-Distance Field (SDF) and Direct Check (DC) evaluations: (a) increasing mesh resolution and (b) increasing number of particles tracked.

## 5 Application to Biomedical Flows

In this section, we apply our SDF-based algorithm to simulate fragmented clot and embolus movement in a patient-specific arterial segment, a representative biomedical simulation involving fluid-particle interactions. Identifying embolus trajectory and distribution is critical for diagnosis and long-term care management of stroke patients, yet current imaging or clinical data cannot provide this information, making *in silico* assessment of embolus movement a promising alternative. In prior works [18, 36], we have established a methodology where an ensemble of embolic particles of varying sizes (*and composition*) was seeded within a patient-specific vascular network model, and their source to destination movement as well as final distribution to the cerebral arteries were computed as important computational descriptors for stroke. In these investigations, we used existing open-source particle tracking software, which implemented a geometric particle contact check with the vessel surface triangulation where points on the sphere were checked for intersection with the triangles along the surface (*that is, no usage of SDF as a means for particle-wall interactions*). Here, we use a representative vascular segment from a set of patient-specific geometries already established in our prior studies [36, 37]. This full segment and corresponding embolus release site is illustrated in Figure 8, panel a comprising the common carotid artery that bifurcates into the external and internal carotid arteries, and subsequently transitions to the cerebral arteries in the brain. This vascular segment was chosen because it is a critical pathway for emboli to enter the brain from the upstream vascular routes from the heart and the aorta. A portion of this segment is shown in Figure 8, panel b alongside the computed 3D SDF field illustrated in Figure 8, panel c. Previously computed CFD simulation data for hemodynamics for this patient specific model was used to impose the background flow velocity through this segment. The hemodynamics data for this was computed using a residual-based stabilized finite element simulation, with custom tuned resistance based boundary conditions to account for the effect of the truncated distal vascular beds on the flow [37]. An ensemble of 1000 representative embolic particles were released at the common carotid artery inlet, each particle with a radius of *R*_*p*_ = 0.25 mm. These particles, representing emboli had a density of 1.1 g*/*cm^3^, while the fluid representing blood was prescribed a density of 1.06 g*/*cm^3^ and a viscosity of 0.04 g*/*(cm · s). Staying consistent with the analyses demonstrated in Section 4, we used the same dynamic equation as in Eq. 7 to compare the particle movement computed using our proposed SDF-based approach, and the previously established geometric approach noted above. Each embolus particle was tracked for 0.83 seconds, representing a single cardiac cycle with anumerical integration time-step of 0.00005 seconds. Representative snapshots of particle positions for successive instants in time (T1-T6) are illustrated in Figure 8, panel d. Embolus particle positions computed using SDF-based approach ***<u>x</u>***_*SDF,i*_ are illustrated in red, while those using the geometric approach ***<u>x</u>***_*geo,i*_ are illustrated in green.

**Figure 8:**
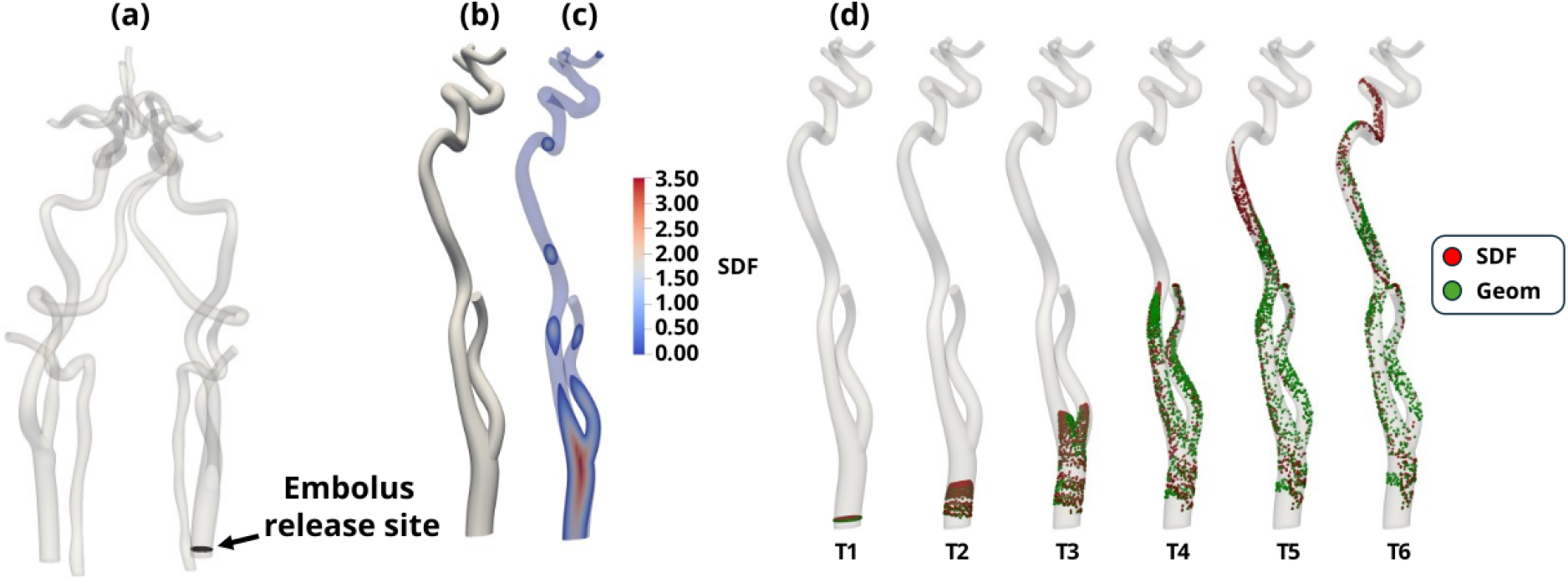
Configuration of a patient-specific vascular model including the whole simulated model with simulated embolus (a), section of the raw geometry (b) alongside the generated SDF field with maximum value in centerline equal to radius of cross-section (c) and sample of particle positions for SDF-algorithm (red) and previously validated particle tracking software (green) tracked particles at 5 time steps (d)

The trajectories generated by the SDF and geometric check implementations were compared using a pair-wise relative difference, where the particles were seeded at identical initial positions and subsequently their trajectories were compared over time. Along their trajectories, the number of collisions that were determined in the SDF algorithm was monitored for each individual particle, in addition to the time instances at which these collisions occurred. A relative difference between the two positions was computed over time as follows:

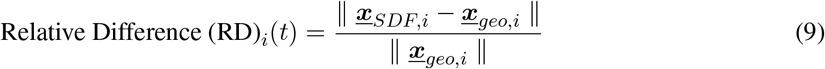

Qualitatively, based on the particle snapshots shown in Figure 8, panel c., it can be observed that the particle positions based on the two algorithms match very well until reaching the first bifurcation in the geometry at T3. It is around this time instant that many particles first collide with the wall. This initial collision occurs early within the full cardiac cycle of tracked motion. Owing to the nature of the dynamics of the particles, small differences between the two algorithms that occur as a result of how collisions are computed subsequently build over time, yielding larger visible differences by the end of the cardiac cycle. The time-averaged relative differences computed over a single cardiac cycle are shown below in Figure 9, panel a. for the entire ensemble of particles. To visualize the effects of the growing differences over time with collisions, the final relative difference compared against the number of collisions is visualized in Figure 9, panel b. The amount of time that particles traveled after their first collision with the wall is also plotted on the graph to provide context as to how much time the previously mentioned small differences have had to accumulate.

**Figure 9:**
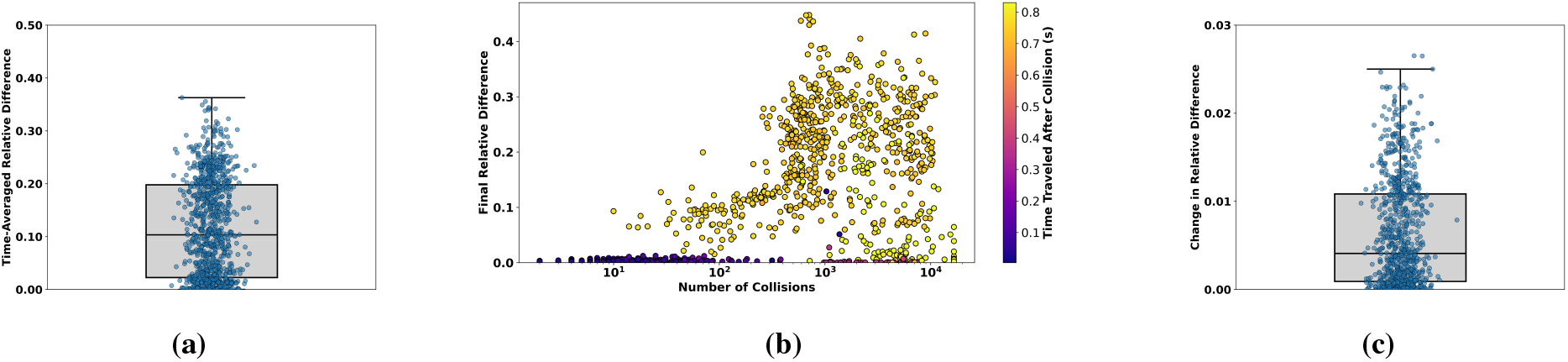
Comparison of relative differences between the SDF-based tracking algorithm and the existing fluid–particle interaction algorithm: (a) time-averaged relative difference (b) final position differences as a function of collision count and post-collision tracking time and (c) change in relative difference after first collision

While we observe that across this ensemble, the mean time averaged relative difference is at 10%, most of the larger discrepancies are associated with particles that undergo frequent wall collisions or are tracked for long periods after their first collision. To further unravel this numerical behavior, the relative difference between the two algorithms was evaluated both before and after the first wall collision. As shown in Figure 9, panel c., a noticeable increase in relative difference (in the order of 2-3% for many particles) often occurs shortly after this initial collision, which, when accumulated over time, contributes to the larger deviations observed in Figure 9, panel a. In addition to comparing the differences in individual particle trajectories, we also illustrate potential differences in aggregate descriptors defined across the trajectory integration, which are commonly used in biomedical contexts. First, we compare the residence time estimates across the two methodologies, indicative of the time taken by the particles to navigate from the entry to one of the multiple outlets of the vascular geometry. This comparison is illustrated in Figure 10, panel a. The mean residence time for the geometric check implementation was 0.64 s, while that obtained from the SDF-based implementation was at 0.57 s. Another quantity of interest for embolic stroke simulations is the distribution of the emboli to the various vessel outlet beds. This is illustrated in Figure 10, panel b. We note that for both aggregate endpoints, there are notable differences between the two wall interaction modeling approaches, as the trajectory differences accumulate over time.

**Figure 10:**
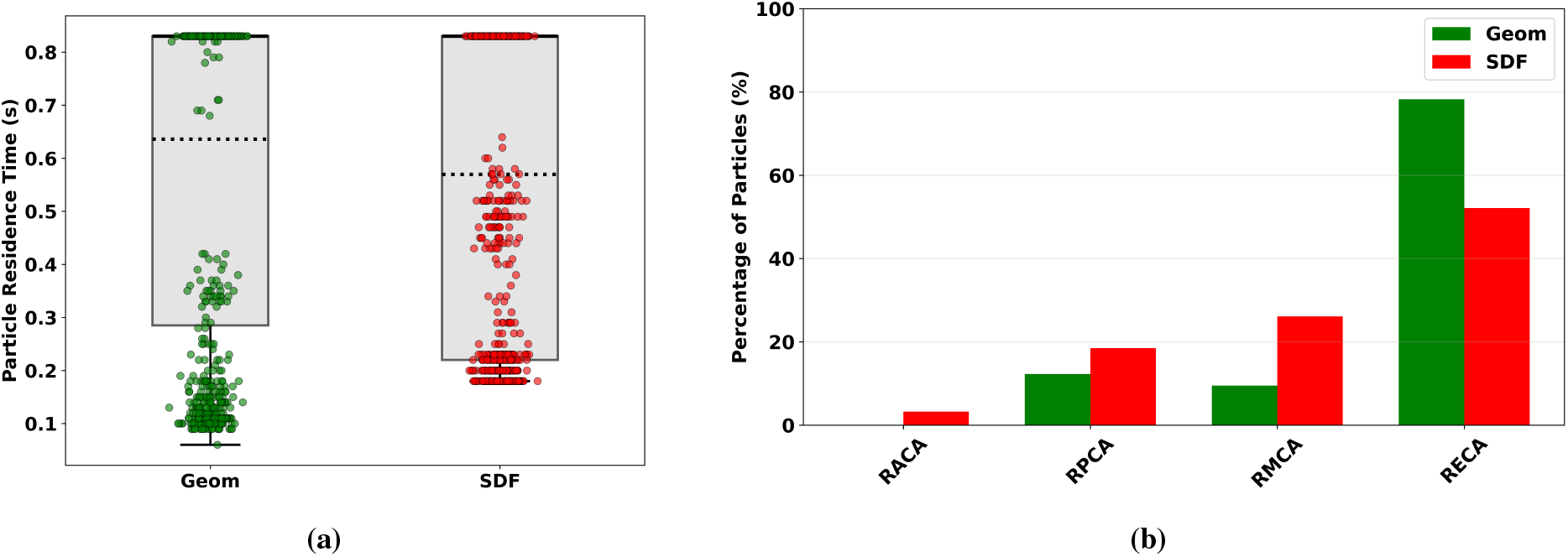
Comparison of derived metrics between the SDF-based tracking algorithm and the existing fluid–particle interaction algorithm: (a) total residence time within the vascular segments and (b) distribution of emboli into each vessel within the vascular network.

These results illustrate few key aspects. First: these two methodologies are independently capable of resolving the nonlinear particle-wall interactions for curved and tortuous geometries representative of anatomical regions of interest, but using different underlying techniques. Second: the resulting trajectories and dynamics estimated using the two approaches, are similar, but not equal, and there are systematic differences originating from underlying methodological features. Third: there is high extent of sensitivity to wall interaction resolution that underlies the dynamics of an inertial particle traversing a nonlinear spatiotemporally varying fluid flow common in biomedical systems. Consequently, the inherent non-linearities, and the underlying dynamical system, can ultimately amplify small errors in particle trajectories accumulated over repeat collisions. These findings together, along with the independently established characterization of accuracy and computational cost in Section 3 and Section 4, help establish the proposed SDF-based approach as a reasonable methodology for complex fluid-particle interaction problems in biomedical flows. This methodology requires that computationally efficient SDF reconstruction is available, and the analyses presented here illustrate that for instances where implicit geometry representation using SDF is not available, conventional geometry based particle-triangle checks are needed to resolve the particle-wall interactions.

## 6 Conclusions

We have developed a particle-wall interaction algorithm for Lagrangian or particle-based computations for physiological and biomedical flow phenomena, using implicit geometry representation based on the Signed Distance Field (SDF). We have demonstrated that the algorithm is computationally efficient. By converting the Lagrangian task of detecting particle-wall proximity to am inherently Eulerian operation, our approach mitigates the increase in computational cost with increasing background and surface mesh resolution typically observed in Lagrangian computations. We have also demonstrated that our approach produces particle trajectories consistent with both analytical examples, as well as conventional wall collision checking approaches. Finally, we have demonstrated that our proposed approach can be efficiently implemented for representative physiological applications involving human subject-specific anatomical features, such as the vascular network in the brain. We remark that the flexibility of implicit geometry representation, and accounting for complexities like curvature and tortuosity, as enabled by our proposed approach is particularly beneficial for physiological and biomedical applications. A direct experimentally derived validation exercise for such an approach was out of the scope given the substantial complexities associated with such experiments. However, we note that further numerical advancements and benchmarking of this method will benefit from such targeted experimental calibration and validation studies.

While the focus of this study was on methodology development and characterization, there are several additional implications that are important to note as key advantages of the proposed implicit geometry based approach. First, given the use of SDF as the methodology for wall proximity and interaction modeling, our approach can essentially be extended to any structure or collection of points not necessarily limited to individual particles. This renders broader applicability of our algorithm for scenarios involving medical devices, stents, and intra-luminal catheters where device or component interaction with vessel or organ wall is of importance. Second, the implicit SDF based representation also holds key advantages in terms of moving mesh or moving boundary scenarios, where there are possible ways to simplify accounting for particle or structure interactions with moving walls or a vessel or organ, based on efficient updates of SDF to represent wall motion. This can enable further computational gains in complex Lagrangian simulations. Third, given that anatomical information is commonly directly obtainable from medical imaging (CT/MR/ultrasound), it remains possible to map out the SDF field and obtain implicit geometry representations directly from images. This can enable rapid device-wall interaction estimates which is important for endovascular or en-doluminal navigability. Additionally, this can also enable direct incorporation of Lagrangian computations with flow-resolving medical imaging such as 4D flow MRI or vector flow ultrasound; thereby substantially accelerating *in silico* interpretation of key descriptors from these images. These broad implications originate specifically from the mathematical and numerical origins of the algorithm as outlined here. While we have not explicitly illustrated these as use cases here, such investigations will remain a focus of future extensions of this methodology.

## Conflicts of Interest

Authors have no conflicts of interest regarding this study and the contents of this manuscript.

## Acknowledgments

This work was partly supported by the National Science Foundation under grant number OAC-1750865 (AK), the National Institutes of Health under grant number R21EB029736 (DM), and the University of Colorado Anschutz-Boulder (AB) Nexus Research Collaboration Award (DM).

## Figures and Tables

### Algorithm 1

Particle Collision

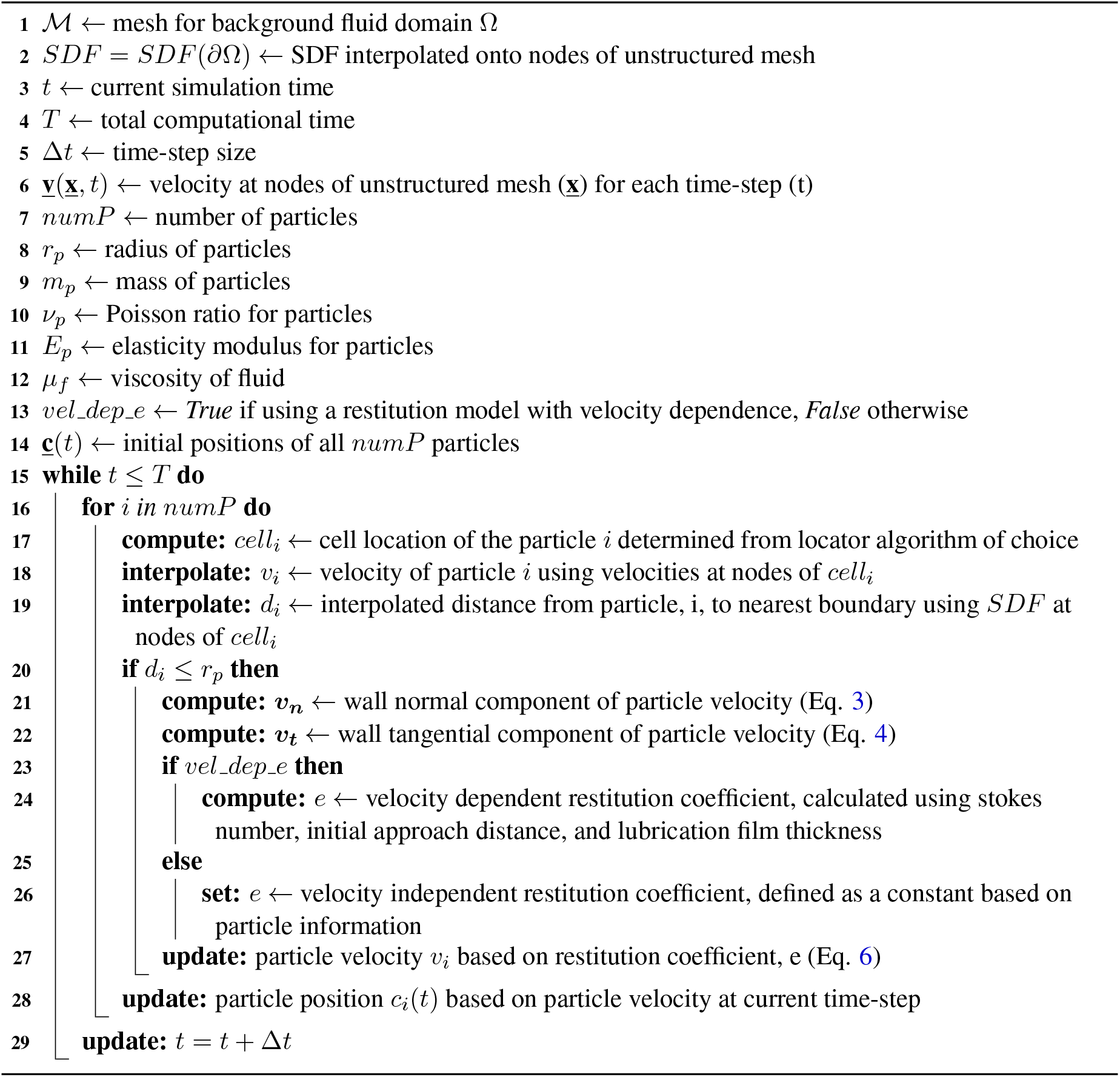

## Notes

### Competing Interest Statement

The authors have declared no competing interest.

